# Detecting the genetic variants associated with key culinary traits in *Dioscorea alata*

**DOI:** 10.1101/2023.10.18.562904

**Authors:** Komivi Dossa, Mahugnon Ezékiel Houngbo, Mathieu Lechaudel, Erick Malédon, Yedomon Ange Bovys Zoclanclounon, Jean-Luc Irep, Mian Faisal Nasir, Hâna Chair, Denis Cornet

## Abstract

Quality attributes play a pivotal role in determining consumers’ acceptability and market value of food crops. *Dioscorea alata* is a major yam species for food security in tropical areas, but our understanding of the genetic factors underlying tuber culinary traits is limited. This study aimed at elucidating the genetic basis of key culinary attributes, including dry matter content, cooking time, boiled yam hardness, and moldability, through genome-wide association studies (GWAS). Phenotypic assessments revealed notable variations among the *D. alata* genotypes across two locations as well as significant correlations among the quality traits. The GWAS identified 25 significant associations distributed across 14 chromosomes. Allele segregation analysis of the identified loci highlighted favorable alleles associated with desired traits, such as reduced cooking time, increased dry matter content, enhanced hardness, and good moldability. Within the set of 42 putative candidate genes, we identified specific genes differentially expressed in tubers of distinct genotypes with contrasting quality attributes. Furthermore, we conducted a comparative analysis with previously reported quantitative trait loci for dry matter content and showed that multiple genomic regions govern this trait in *D. alata*. Our study offers valuable insights into the links between these key culinary traits and the underlying genetic basis in *D. alata*. These findings have practical implications for breeding programs aimed at enhancing the quality attributes of greater yam.

## Introduction

Yams (*Dioscorea spp*.) are sources of essential carbohydrates in various regions of the world, holding cultural and economic importance, particularly in West Africa [1–3]. While a few Dioscorea species are used for food and industrial purposes, the most extensively cultivated ones include *D. alata*, *D. cayennensis*, and *D. rotundata* [4]. *Dioscorea rotundata* is indigenous to West Africa and has the largest production volume, while *D. alata* was introduced to Africa from Asia and is the most commonly grown species globally [5]. Traditional yam breeding has long overlooked quality traits, as phenotypic evaluation of these complex properties is slow and labor-intensive [6]. However, emerging tools now empower more efficient dissection of the culinary quality (https://rtbfoods.cirad.fr/). Recent advancements in near-infrared spectroscopy have shown promise in rapidly predicting quality traits such as texture and biochemical composition in yams, which can significantly accelerate breeding programs and enhance food security [7–9].

Cooking quality is a crucial aspect of yam acceptance by consumers [10, 11]. Traits like hardness and moldability (Figure 1) are essential factors considered by consumers and processors when selecting yam varieties for cooking and processing purposes [12]. Hardness refers to the firmness or resistance of the cooked yam flesh. It is an important quality attribute as it affects the sensory experience and ease of consumption [13, 14]. Hardness is influenced by factors such as the composition of cell walls, starch structure, and secondary metabolites [15]. For instance, a higher proportion of amylose, a starch component, is associated with increased hardness in sweet potato [16]. On the other hand, softer yam varieties typically have a higher moisture content and lower starch content. The cooking method, such as boiling or steaming, also affects the final hardness of yam [17].

**Figure 1.**
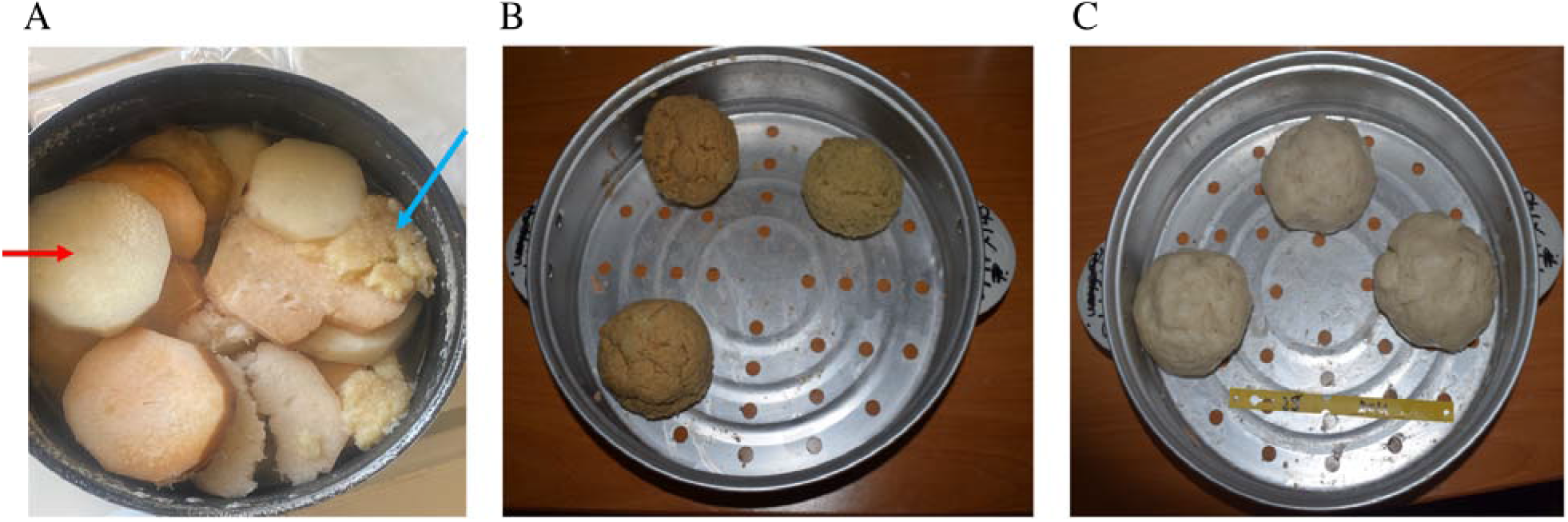
Illustrations of boiled and pounded *D. alata* tuber. **A**) Tubers of various genotypes were boiled, and different hardness levels could be observed. The blue arrow shows an example of a genotype with a low hardness, while the red arrow shows a variety with a high hardness. **B**) A genotype with a bad moldability quality; **C**) A genotype with good moldability quality.

Moldability, or poundability, refers to the ability of cooked yam to be pounded or mashed into a desired consistency [7, 18]. It is a desirable trait for traditional dishes like pounded yam in West Africa, where yam is traditionally pounded with a mortar and pestle [19]. Cassava varieties with higher starch content, particularly varieties with a higher proportion of amylopectin, tend to exhibit better moldability due to the retrogradation of starch [20].

Cooking time is an important quality trait in yam (*Dioscorea spp*.), influencing consumer’s acceptance [21]. Depending on species and variety, it can range from 10 minutes to over a hour, while short cooking time is preferred [21]. Although the correlation between cooking time and dry matter content is not established in yam, other tubers, such as cassava (*Manihot esculenta*) and Japanese taro (*Colocasia esculenta*), tend to show a positive correlation between them [22–24]. Based on these reports, it can be assumed that dry matter content, representing starch and fiber levels, is positively associated with longer cooking time.

Efforts to enhance the cooking quality of yam, particularly hardness and moldability, in breeding programs involve genetic studies [25–28]. *Dioscorea alata* is renowned for its favorable agronomic performance and widespread cultivation. It exhibits a stable yield, making it well-suited for addressing the challenges posed by the current climate change scenario [29–31]. However, one significant drawback of this crop is its relatively low culinary quality [32]. In contrast, *D. rotundata* and *D. cayenensis* are characterized by superior culinary attributes. Unfortunately, interbreeding between these species is challenging due to cross-incompatibility [33]. Within *D. alata*, there has been observed a diversity in culinary quality, as documented in studies like Ehounou et al. [7, 34]. Consequently, further genetic investigations focusing on the underlying genetic factors governing culinary quality traits in *D. alata* are necessary.

The availability of a high-quality genome sequence for *D. alata* has opened a new avenue for genetic and genomic studies of tuber quality traits [35–39]. Transcriptome profiling, quaantitaive trait locus (QTL) mapping, and genome-wide association studies (GWAS) have identified genomic regions and candidate genes associated with various yam quality attributes, such as tuber color, texture, dry matter content, and starch accumulation [5, 25, 36, 40–42]. These findings pave the way for future efforts in genetic engineering and breeding programs to develop superior yam varieties with improved nutritional and sensory qualities. Integrating genomics and breeding approaches is crucial for enhancing yam production, ensuring higher yields, better taste, and increased nutritional value, ultimately benefiting communities worldwide that rely on yam as a staple crop.

In this study, the main objective was to unravel the genetic basis underlying the culinary quality parameters, including cooking time, dry matter content, hardness, and moldability in *D. alata*. To achieve this, a diverse panel of *D. alata* accessions was carefully selected, and these accessions were evaluated for the desired traits at two locations. The combination of GWAS and gene expression profiling was employed to identify the specific genomic regions and potential genes associated with the culinary quality traits.

### Plant material and experimental design

The plant material utilized in this study consisted of a diverse panel of *D. alata* accessions [43]. The panel comprised 52 diploid accessions collected from various tropical countries, including Africa, the Caribbean, and the Pacific. The field trials were fully detailed in Dossa et al. [43].

### Assessment of cooking time

Following the field harvest of the 52 genotypes, three tubers per genotype (biological replicates) were selected. To cook the tubers, three identical pressure cookers (Moulinex) with 3L capacity and basic kitchen utensils were employed. The method described by Ehounou et al. [7] was followed. First, the head and the tail were removed. Then the tubers were peeled, sliced (∼ 1 cm thickness), and cooked in parallel with 1.5 L water in the pressure cookers until they reached the desired level of softness, determined using the fork test. The cooking time (min) was recorded subsequently for each tuber.

### Assessment of moldability

Moldability is the ability of the dough from the pounded yam to form a ball. Three mechanical pounders (Moulinex) were used for this experiment following the descriptions from Ehounou et al. [7]. Although the dough obtained from a mechanical pounder is not exactly the same as the one obtained with the traditional practice of manual kneading in a mortar with a pestle, our method ensures uniform conditions for sample preparation and guarantees the repeatability of the experiment. Once the boiled yams were ready, 10 slices from each tuber were randomly selected and inserted into the mechanical pounders. Each pounding lasts for 2 min, and the dough was assessed for its moldability quality based on a score within 0-9, with 0 meaning no ball could be formed in hand while 9 represents a ball is perfectly formed, and the overall texture is very similar to the best-pounded yam obtained from the traditional practice.

### Assessment of dry matter content

In addition to the cooking process, the remaining portion of the yam samples that were not used for cooking underwent further analysis. These samples were cut into small pieces, weighed, and then dried in an oven for 72 h at a temperature of 60°C. After drying, the samples were weighed again to determine their dry weight. The dry matter content of the samples was calculated using the following formula:

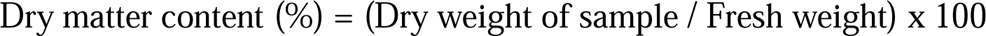

The dry matter content is expressed as a percentage and provides valuable information about the proportion of solid matter in the yam samples after moisture removal through drying.

### Assessment of hardness

To estimate the hardness, Texture Profile Analysis (TPA) was conducted in this study as previously implemented by Mota et. al. [37]. The TPA measurements were performed at 45°C using the TAX-T2 texture analyzer from Stable Micro Systems. The experiment employed a combination of Surrey Limited hardware and Exponent software to facilitate data capture, storage, and real-time analysis. For measurement, three tubers per genotype were utilized. Only the central part of the tuber was selected for analysis in order to maintain consistency and standardization throughout the experiment. From the central part of each tuber, three 23 mm cubes were cut and steam-cooked in a pressure cooking for 15 min. Before measuring, the cooked cubes were cooled for 7 min. Hardness is expressed in N/cm² and defined as the strength needed for the cube to be compressed and deformed. TPA measurements were made on tubers from Godet only. Due to limited resources, tubers from Roujol could not be processed on time for a valid comparative analysis.

### Genome wide association studies and characterization of alleles

Genotyping data from a panel of 52 diverse genotypes was obtained from our previous study [36]. A genome-wide association study (GWAS) was conducted using 1.9M high-quality single nucleotide polymorphisms (SNPs), and 45,000 randomly selected SNPs from the initial 1.9M datasets. The two SNP densities were tested to highlight the effect of SNP density on GWAS power. The SNP density plot was generated using the “CMplot” package [44] in R4.0.23. GWAS was performed using two statistical models: Fixed and random model circulating probability unification (FarmCPU) and mix linear model. The FarmCPU algorithm has enhanced statistical power and computational efficiency and can mitigate false associations in GWAS [36]. To identify SNPs significantly associated with traits in each environment, the adjusted p-value was utilized. A significance threshold of P < 10^-8^ (0.05/n, with n being the number of SNPs) was employed to identify significant associations. Additionally, quantile-quantile (QQ) plots were constructed to assess the extent to which the models accounted for population structure by comparing the observed p-values to the expected p-values under the null hypothesis of no association.

### Statistical analysis

The impact of alleles at significant SNPs was evaluated by comparing phenotyping data among different genotypic groups. Student’s t-test, with a significance level of P < 0.05, was conducted using XLStat 2023.5.1.1399 to compare the genotypic groups.

To analyze genotype by environment interactions, a two-way analysis of variance (ANOVA) was performed using the statistical software Statistics Analysis System (SAS 9.4, SAS Institute Inc., with genotype and environment as the main effects. A significant genotype x environment interaction indicated differential phenotypic response across environments. Broad sense heritability was estimated by obtaining variance components through REML in ASReml and taking the ratio of genetic variance to the total phenotypic variance. Pearson’s correlation coefficients between trait pairs were calculated using standard functions in SAS 9.4, SAS Institute Inc. Together, these analyses enable the characterization of genotype-by-environment effects, total genetic contribution to traits, and trait correlations to elucidate the genomic factors underlying complex phenotypic patterns.

### Comparative *in silico* analysis of quantitative trait loci for dry matter content in *D. alata*

To construct a physical map of quantitative trait loci (QTL) for DM in *D. alata*, marker sequences were obtained from two previous QTL mapping studies by Bredeson et al. [35] and Gatarira et al. [41]. The marker sequences were then aligned to the *D. alata* V2 reference genome assembly [35] using BLAST to determine the specific physical positions of the reported QTL regions. Additionally, quantitative trait nucleotides (QTNs) identified from GWAS in the current study were mapped to precise genome locations. All collected QTL positions and QTNs associated with DM were integrated into a comprehensive physical map using Mapchart version 2.32. This consolidated physical map provides a genomic framework for further characterization of sequence variants and candidate genes influencing DM in *D. alata*.

### Expression profiling of candidate genes

Transcriptomic data from tubers of three contrasting genotypes, including Roujol49, Roujol75, and Roujol9, with three biological replicates [37], were utilized to characterize the putative candidate genes based on their corresponding expression profiles. The genotypes were selected, based on their culinary qualities and corresponding differential alleles at the peak SNPs, to further investigate the comparative transcriptomic profile and identify putative candidate genes related to each trait under study. A gene was considered differentially expressed if there was a statistically significant difference with a threshold of log_2_ fold change |log_2_FC| ≥ 1, false discovery rate adjusted P-value (padj) ≤ 0.05 in pairwise comparisons.

## Results

### Phenotypic characterization of the diversity panel

Table S1 presents data on 52 genotypes of yams and their corresponding measurements for traits such as dry matter content (DM), cooking time, moldability, and hardness (Figure 2 and Table S1). The mean values for these traits were 29.12% for DM, 18.50 min for cooking time, 5.30 for moldability, and 28.22 for hardness. The range of values varied across the genotypes, with the minimum and maximum values being 20.55% and 37.28% for DM, 11 min and 33 min for cooking time, 1.67 and 9 for moldability, and 6.39 and 85.73 for hardness. The coefficient of variation (CV) indicated the degree of variation within the traits, with CV values ranging from 10.54% to 60.50%. These results provide valuable insights into the diversity of yam genotypes in relation to their cooking quality, which can be harnessed in breeding programs.

**Figure 2.**
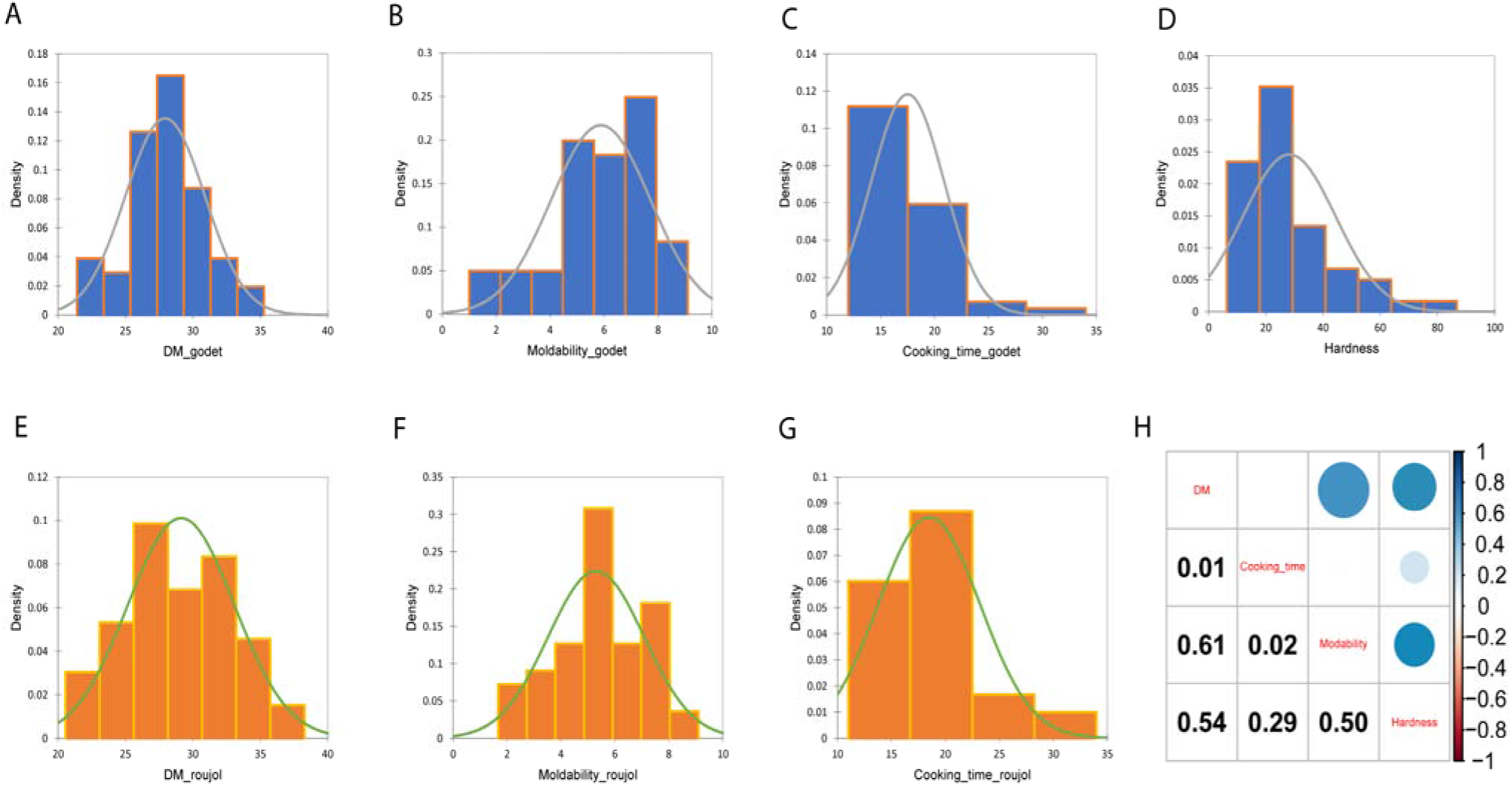
Phenotypic variation estimated for traits under study. **A)** Dry matter content (DM) at Godet, **B**) Moldability at Godet, **C)** Cooking time at Godet, **D)** Hardness at Godet, **E)** DM at Roujol, **F)** Moldability at Roujol, **G)** Cooking time at Roujol, and **H)** Correlation between the different traits under study.

Overall, the observed differences in traits between Roujol and Godet indicate the environmental effects on the culinary characteristics of *D. alata* (Table 1). These differences could be attributed to variations in environmental conditions, soil composition, or other factors specific to each location.

**Table 1.**
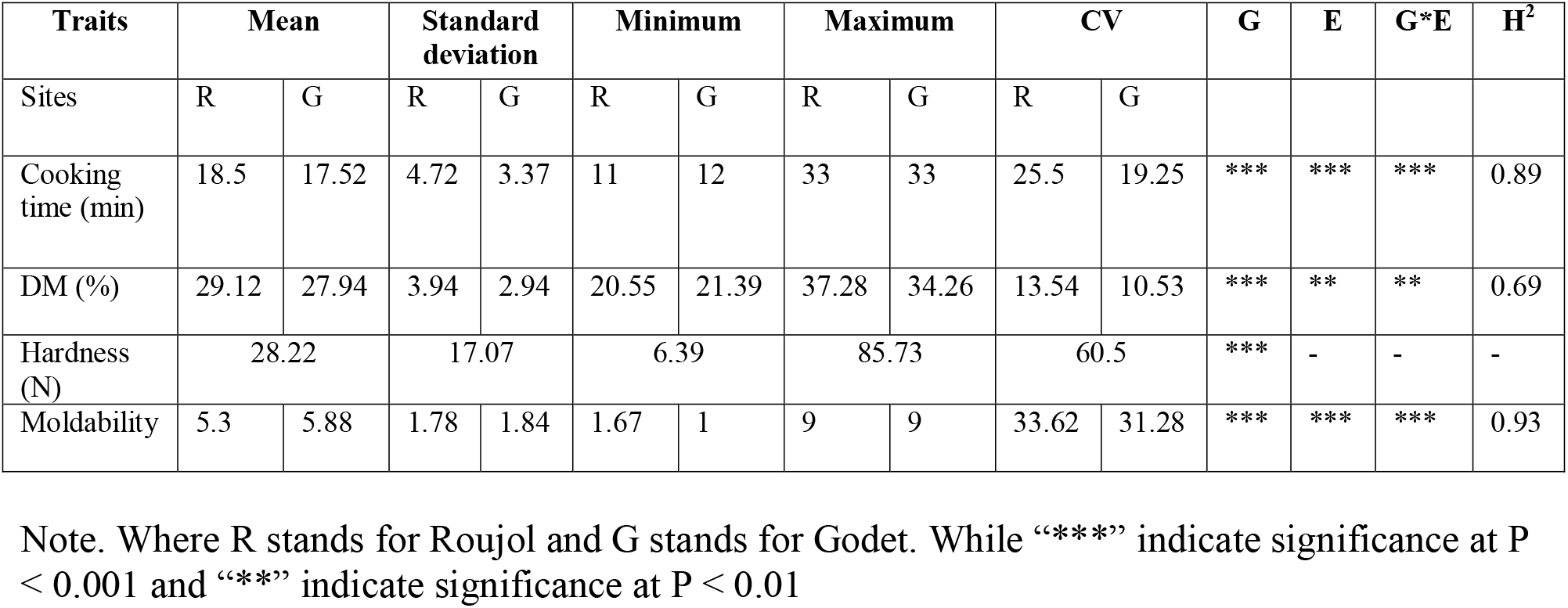
Statistics of the phenotypic traits among the varieties and planting sites.

We further estimated the correlation coefficients between the traits (Figure 2H). The correlation analysis for yam traits reveals that DM is strongly and positively correlated with moldability and hardness. This observation means *D. alata* varieties with higher dry matter content exhibit better moldability and higher hardness. Cooking time shows a weaker correlation with moldability and hardness. Moldability and hardness in yam are strongly and positively correlated, indicating that boiled yam hardness can be used to predict moldability quality. Moreover, cooking time and moldability depicted high heritability, 0.89 and 0.93, respectively, implying that breeding for these traits in *D. alata* will be amenable.

### Impact of SNP density and statistical model on GWAS result

In our previous study, we performed a comprehensive genetic analysis on our diversity panel consisting of multiple genotypes and showed a low structure and its suitability for GWAS [36]. In order to determine how statistical models and SNP density affect GWAS, we selected two traits, cooking time and DM, and performed GWAS with two SNP densities: low density with 45,000 SNPs (Figure 3A, 0.09 SNPs/kb) and high density with 1.9M SNPs (3.9 SNPs/kb)) as well as two statistical models (FarmCPU and MLM). The results showed that high-density SNP data improves GWAS power with highly significant associations (-log_10_P > 9), as depicted in Figures 3B and 3C. Additionally, the FarmCPU model outperformed the MLM model with more significant associations (-log_10_P > 7.5), as shown in Figures 3D and 3E. These findings suggest that utilizing high-density SNP data and employing the FarmCPU model enhances the accuracy and power of GWAS in *D. alata*. The 1.9M SNPs dataset was thus kept for downstream analyses.

**Figure 3.**
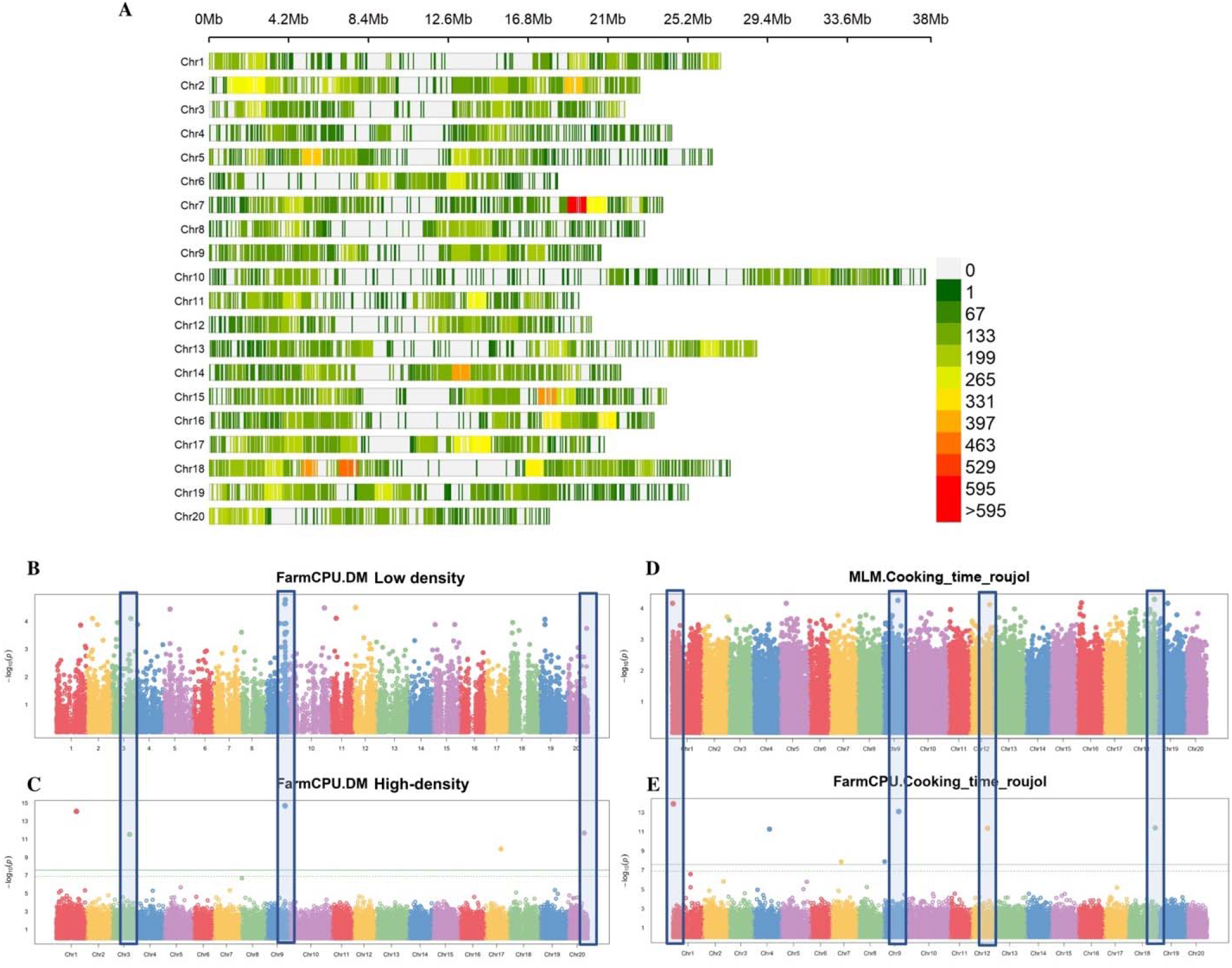
Effect of SNP density and model selection on GWAS. **A)** subplot A displays the SNP density at a lower level, with 45,000 SNPs. The plot with 1.9 M SNPs was presented by Dossa et al. [36]. This plot illustrates the distribution and density of SNPs across the genome, indicating the coverage and resolution of the genotyping data used in the GWAS analysis **B)** The Manhattan plot in subplot B represents the GWAS associations obtained using the lower SNP density. The Manhattan plot shows the genomic positions of SNPs on the x-axis, with their corresponding association p-values on the y-axis. The peaks in the plot indicate significant associations between SNPs and the studied trait. The lower SNP density may result in fewer significant associations being detected**, C)** Manhattan plot showing GWAS associations with high SNP density. This plot represents the GWAS using a higher density of SNPs, which improves the coverage and resolution of the genotyping data. With higher SNP density, more potential associations between SNPs and the trait can be identified, **D)** Manhattan plot showing GWAS associations estimated using mixed linear model (MLM) for cooking time at Roujol, **E)** Manhattan plot showing GWAS associations estimated using FarmCPU for cooking time at Roujol.

### GWAS for dry matter content

GWAS for tuber DM identified six significant associations (Log_10_P > 9) at the two locations, Roujol and Godet (Table 2). In Roujol, significant associations were found on chromosomes 1, 3, 9, 17, and 20 (Figure 4A, 4B), while in Godet, a significant association was observed on chromosome 9 (Figure 4C, 4D). Interestingly, the genomic region on chromosome 9 was consistently identified in both locations. The variation in GWAS results between the two locations could be attributed to environmental factors impacting the phenotype. To gain further insights into the effects of the identified loci, allelic effects for the top significant or stable SNPs (rs839254 and rs50615) were investigated using allele segregation analysis.

**Figure 4.**
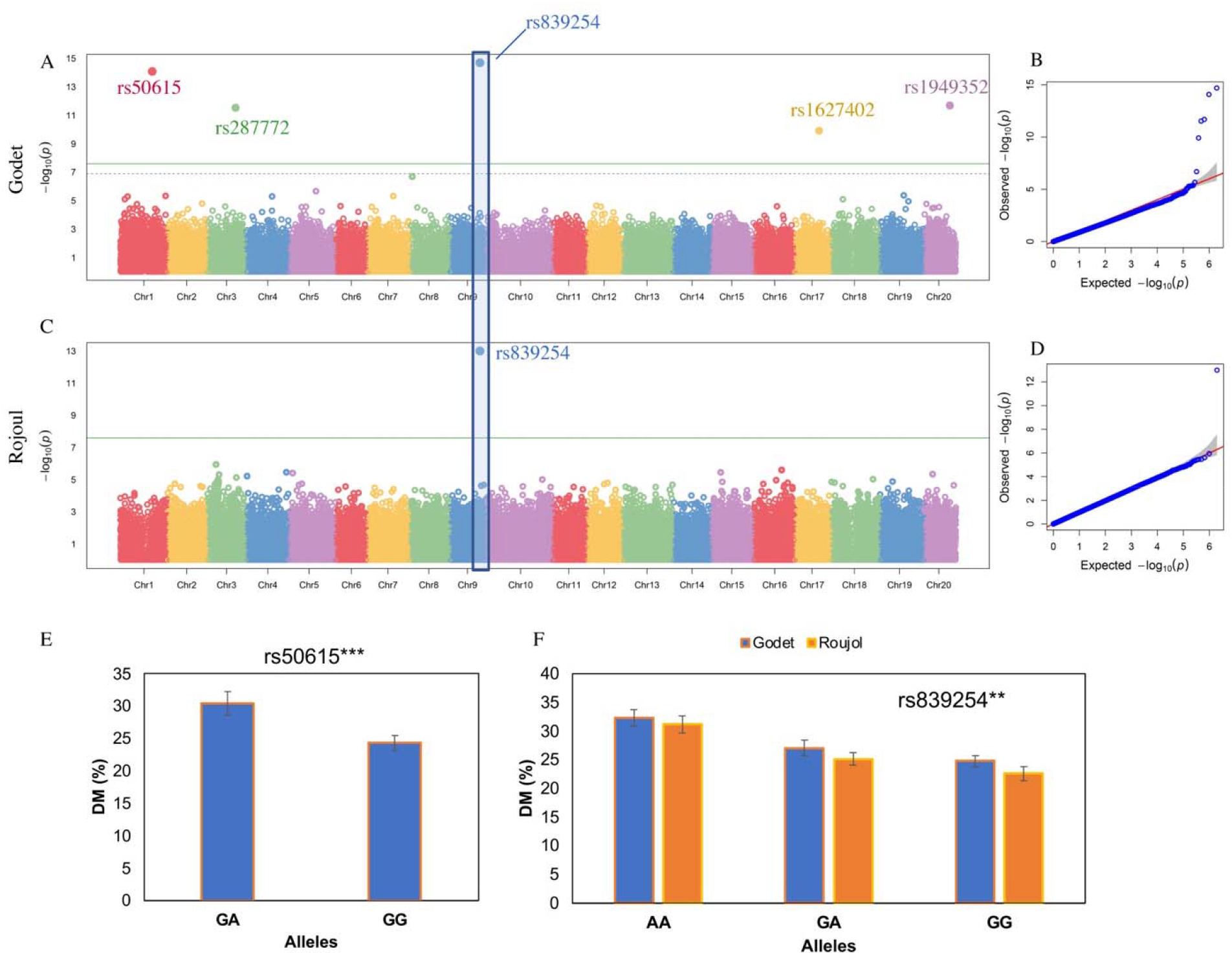
GWAS for dry matter content (DM). **A)** The Manhattan plot for DM at Godet shows the genomic positions of SNPs plotted against their association p-values. The peaks in the plot indicate significant GWAS signals. The green horizontal lines represent the genome-wide significance threshold, helping to identify SNPs that surpass the threshold and are considered highly associated with DM, **B)** The QQ-plot associated with DM at Godet depicts the observed p-values against the expected p-values under the null hypothesis of no association. Deviations from the diagonal line suggest potential associations between SNPs and DM, **C)** The Manhattan plot for DM at Roujol displays the genomic positions of SNPs and their association p-values specifically for the DM at Roujol, **D)** The QQ-plot associated with DM at Roujol is similar to the plot described in (B), **E)** The allele segregation analysis concerning SNP rs50615 demonstrates the relationship between different genotypes and their corresponding cooking times. In this analysis, the GA allele is associated with genotypes that have higher cooking time, while the GG allele is associated with genotypes that have lower DM, **F)** The allele segregation analysis concerning SNP rs839254 demonstrates the relationship between different genotypes and their corresponding cooking times. In this analysis, the GA allele is associated with genotypes that have medium DM, and the AA allele is associated with genotypes that have higher DM, while the GG allele is associated with genotypes that have lower DM. ** indicates significant differences at *P* = 0.01 and *** corresponds to significant difference at P = 0.0001

**Table 2.**
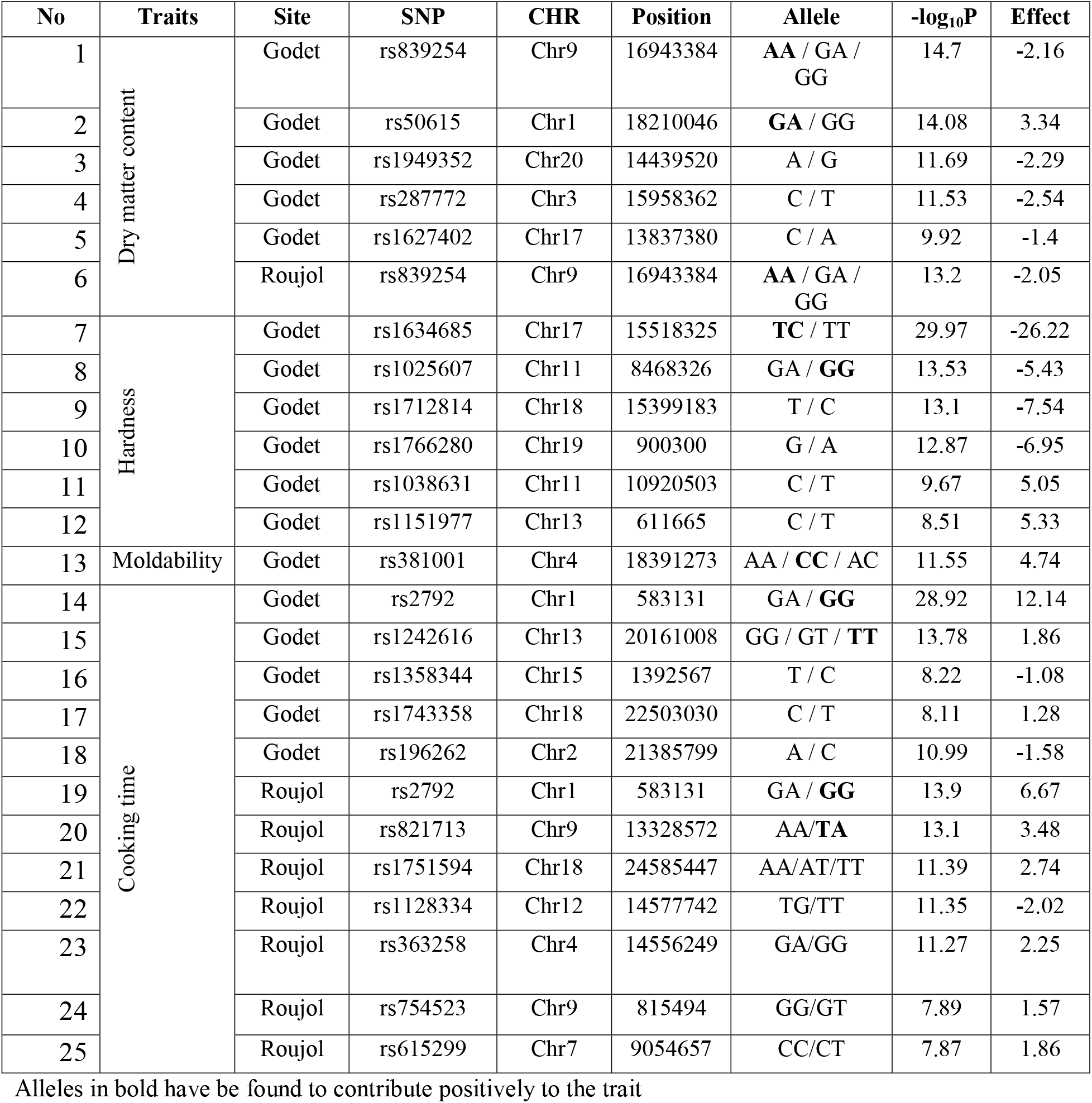
The significant signals identified with the allelic effects for each SNP.

The SNP rs839254 consistently appeared in the same genomic region on chromosome 9 at both locations (Table 2). It exhibited clear differentiation of genotypes into three allelic groups: AA, GA, and GG. Homozygous accessions (AA) at this locus were associated with higher DM compared to AG and GG genotypes (Figure 4F). SNP rs839254 is annotated as a downstream SNP of *Dioal.09G047900*, which encodes a Winged helix-turn-helix DNA-binding domain protein (Table 3).

**Table 3.**
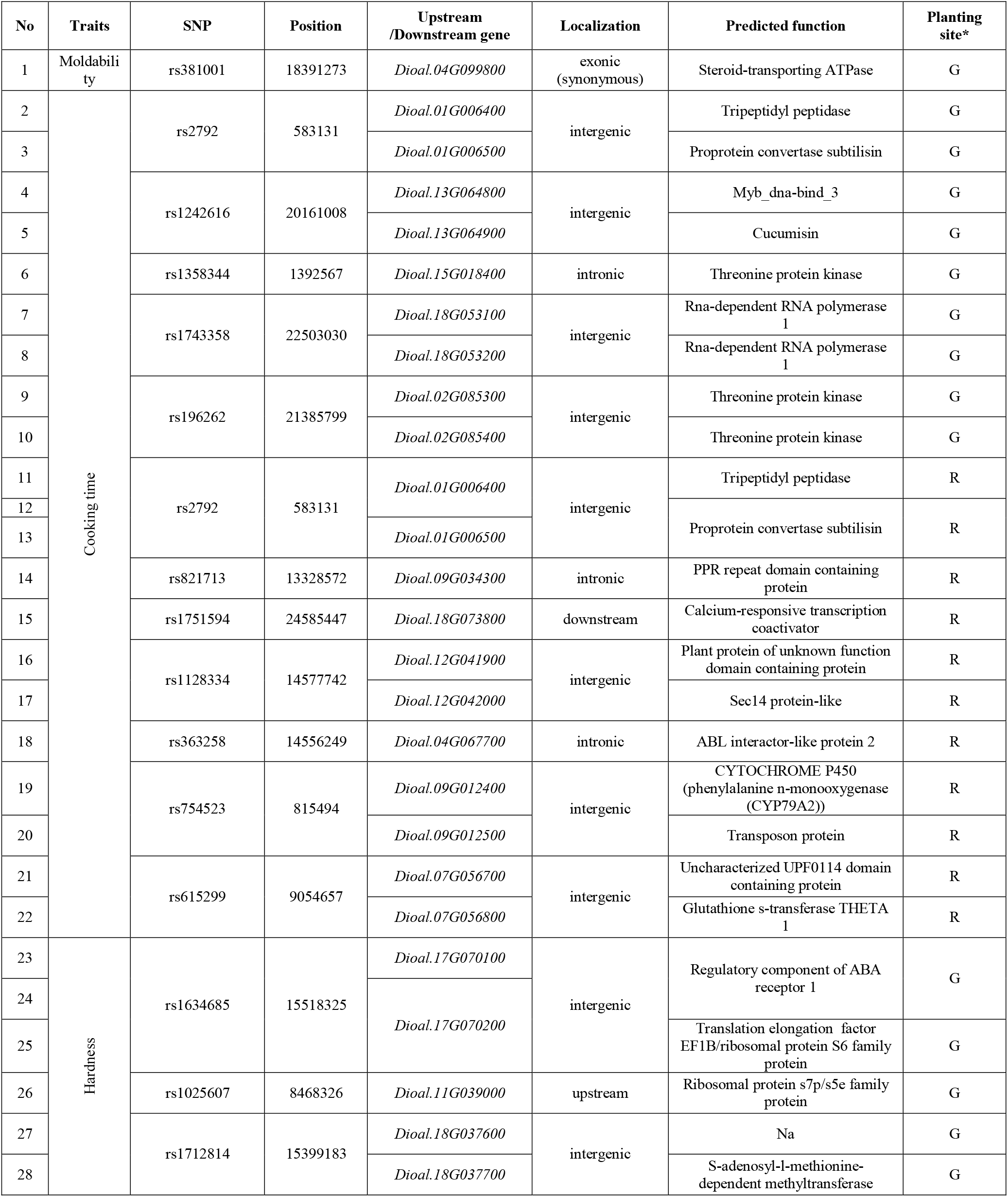
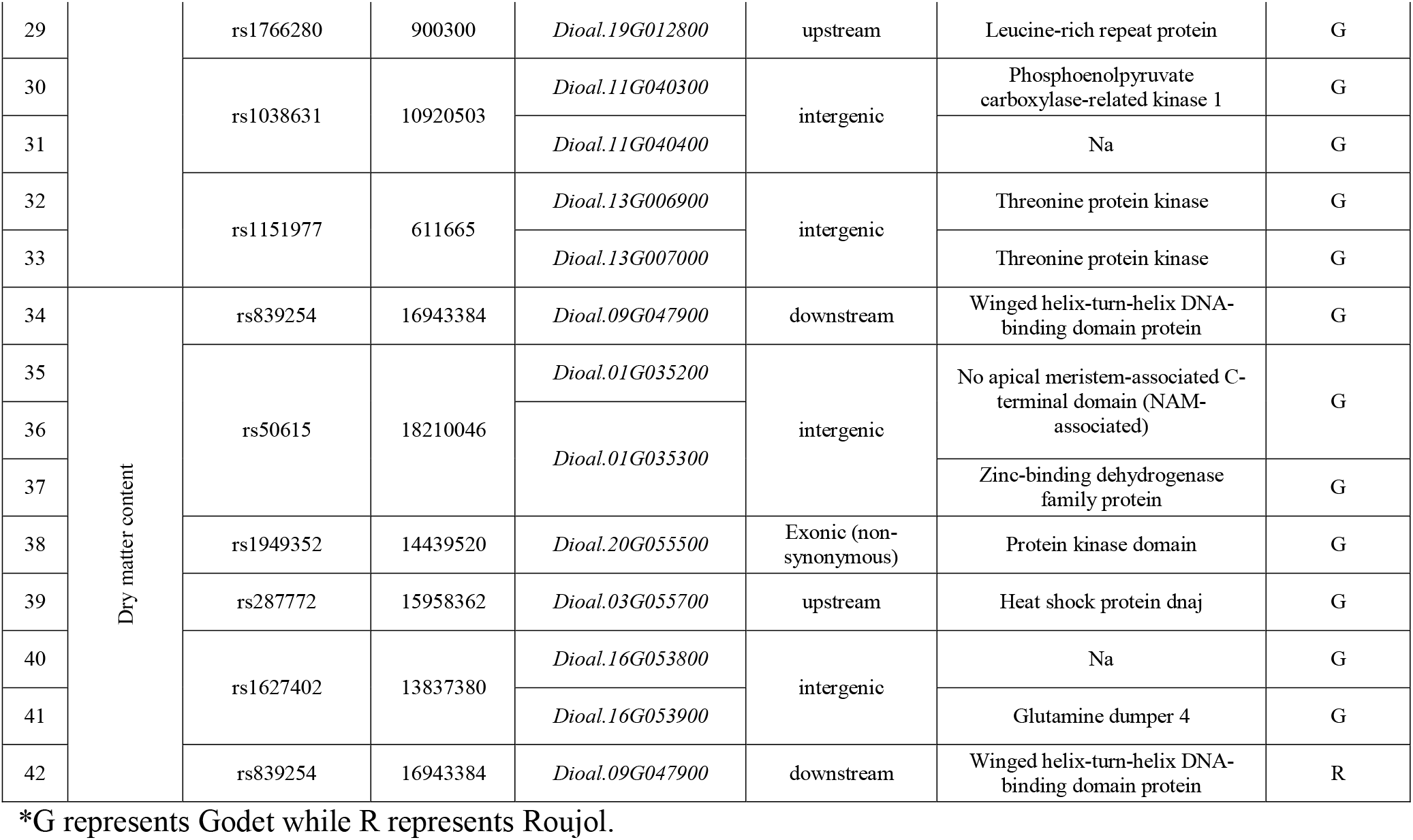
Summary of identified candidate genes associated with tuber quality traits in *Dioscorea alata*.

Additionally, we characterized another significant SNP, rs50615 (Log_10_P = 14.08), specifically identified at the Godet location (Figure 4A). Notably, both genotypes GA and GG exhibited significant differences in DM (Figure 4E), with the GA genotype associated with higher DM. The SNP rs50615 is annotated as an intergenic SNP located between the genes *Dioal.01G035200* (No apical meristem-associated C-terminal domain (NAM-associated)) and *Dioal.01G035300* (Zinc-binding dehydrogenase family protein) (Table 3).

### GWAS for cooking time

The GWAS for cooking time revealed eleven significant associations (Log_10_P > 7) at the Roujol and Godet locations (Table 2). In Roujol, significant associations were observed on chromosomes 1, 3, 13, 15, and 14 (Figure 5A, 5B), while in Godet, significant associations were found on chromosomes 1, 4, 7, 9, 12, and 18 (Figure 5C, 5D). One genomic region on chromosome 1 showed consistent associations in both locations. To explore the effects of these identified loci, we performed an allele segregation analysis on the top significant or stable SNP, rs2792.

**Figure 5.**
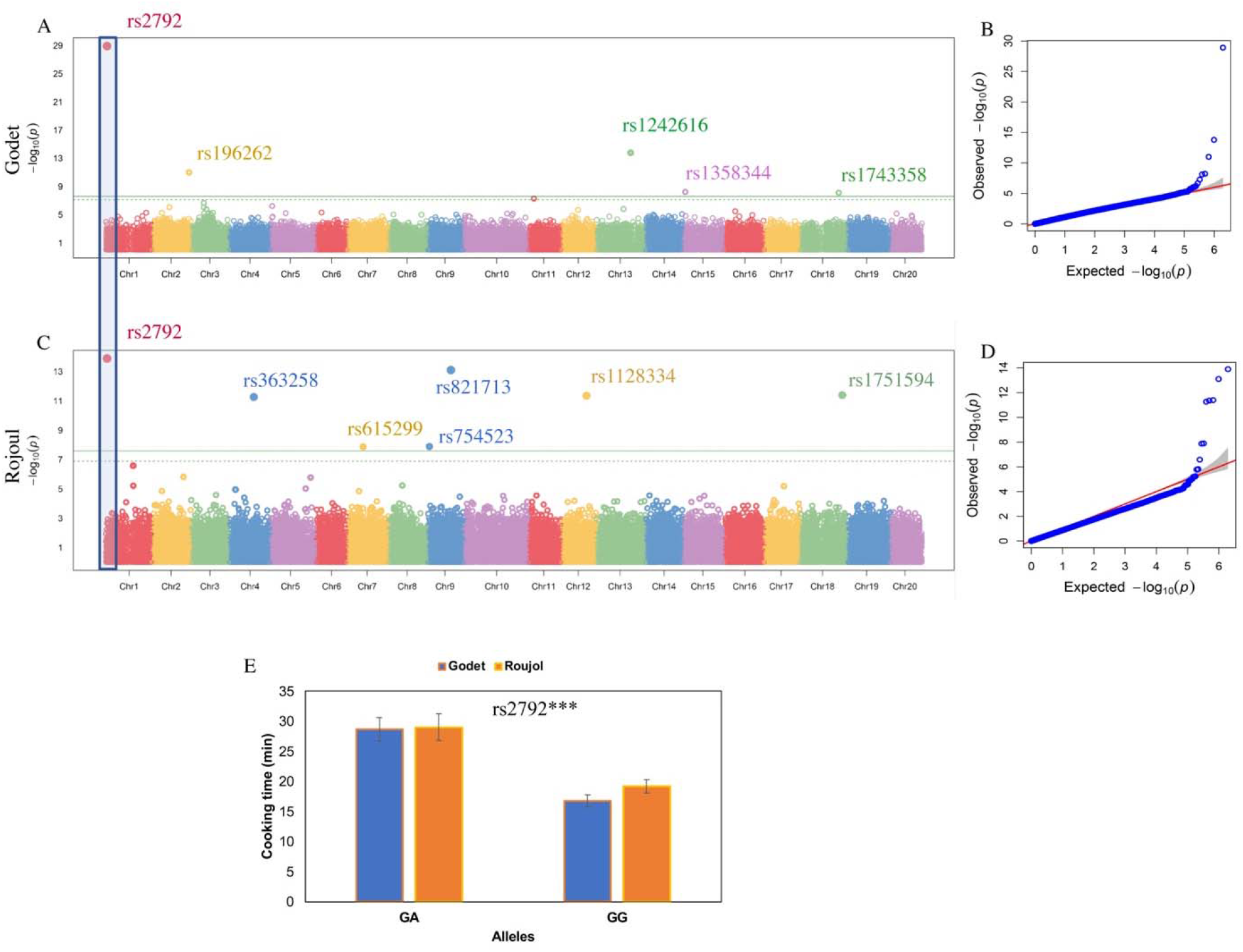
GWAS for cooking time in *D. alata*. **A)** The Manhattan plot for the cooking time at Godet shows the genomic positions of SNPs plotted against their association p-values. The peaks in the plot indicate significant GWAS signals. The green horizontal lines represent the genome-wide significance threshold, helping to identify SNPs that surpass the threshold and are considered highly associated with cooking time, **B)** The QQ-plot associated with cooking time at Godet depicts the observed p-values against the expected p-values under the null hypothesis of no association. Deviations from the diagonal line suggest potential associations between SNPs and cooking time, **C)** The Manhattan plot for cooking time at Roujol displays the genomic positions of SNPs and their association p-values specifically for the cooking time at Roujol, **D)** The QQ-plot associated with cooking time at Roujol is similar to the plot described in (B), **E)** The allele segregation analysis concerning SNP rs176324 demonstrates the relationship between different genotypes and their corresponding cooking times. In this analysis, the GA allele is associated with genotypes that have long cooking times, while the GG allele is associated with genotypes that have short cooking times. *** indicate t significant difference at *P* = 0.0001

The significant SNP rs2792 displayed distinct genotypic differences, with two alleles identified: GA and GG. Accessions with heterozygous genotypes (GA allele) at this locus exhibited longer cooking time compared to those with GG allele (Figure 5E). The SNP rs2792 is annotated as an intergenic SNP located between the genes *Dioal.01G006400* (Tripeptidyl peptidase) and *Dioal.01G006500* (Proprotein convertase subtilisin) (Table 3).

### GWAS for hardness and moldability

For hardness, several significant associations were identified. The SNPs rs1634685 on Chr17, rs1025607 on Chr11, rs1712814 on Chr18, rs1766280 on Chr19, rs1038631 on Chr11, and rs1151977 on Chr13 exhibited strong associations with hardness (Figure S1C). These SNPs had high effect sizes, indicating their influence in determining the hardness of the boiled yam. Allele segregation analysis further confirmed the differentiation of genotypes into two groups based on the TC and TT alleles at SNP;rs1634685 (Figure S2A). Moreover, several candidate genes associated with these SNPs were identified as *Dioal.17G070100, Dioal.17G070200, Dioal.11G03900, Dioal.18G037600, Dioal.18G037700, Dioal.19G012800, Dioal.11G040300, Dioal.11G040400, Dioal.13G006900,* and *Dioal.13G007000* (Table 3).

In the case of moldability, the SNP rs381001 on Chr4 showed a significant association (Figure S1A and S1B). This SNP had three different alleles (AA, CC, AC), and individuals with the AC allele had medium moldability scores (Figure S2B). The effect size for this SNP was positive, indicating that the AC allele contributes to increased moldability. Furthermore, rs381001 was identified as a non-synonymous SNP in the exonic region of *Dioal.04G099800*, annotated as a Steroid-transporting ATPase (Table 3). Further investigations are needed to determine the precise role of this gene in relation to moldability.

### *In silico* comparative analysis of genomic regions identified from previous studies on dry matter

To gain a comprehensive understanding of the genetic control of the DM trait, we compared the identified quantitative trait loci (QTL) from our study with those reported in two previous studies [35, 41]. Surprisingly, we found no overlap between the QTLs identified in the three studies (Figure S3). Although some of the parental genotypes used in the study by Bredeson et al. [35] were also present in the study by Gatarira et al. [41], none of the QTLs were found to coincide. This finding further emphasizes the complex nature of DM, which appears to be influenced by multiple genomic regions and potentially affected by interactions between genotypes and the environment.

### Expression profiling of candidate genes

Transcriptome data [37] of multiple allele-specific genotypes with contrasting phenotypes were analyzed to explore the expression profile of the putative genes detected around the peak SNPs. Out of the 42 candidate genes identified near the peak SNPs on different chromosomes, a subset of 3 and 2 genes showed differential expression pattern for cooking time and dry matter contents, respectively (Figure 6A-6B). *Dioal.02G085400* (rs196262), *Dioal.04G067700* (rs363258), and *Dioal.09G012500* (rs754523), identified as candidate genes for cooking time, showed differential expression (Upregulated in Roujol49 with higher cooking time (32 min) compared to Roujol9 with lower cooking time (15.33 min)). *Dioal.02G085400*, *Dioal.04G067700*, and *Dioal.09G012500* encode Threonine protein kinase, ABL interactor-like protein 2, and Transposon protein, respectively (Table 3). It can be speculated from the results that higher expression of these putative candidate genes result in physiological changes leading to longer cooking time.

**Figure 6.**
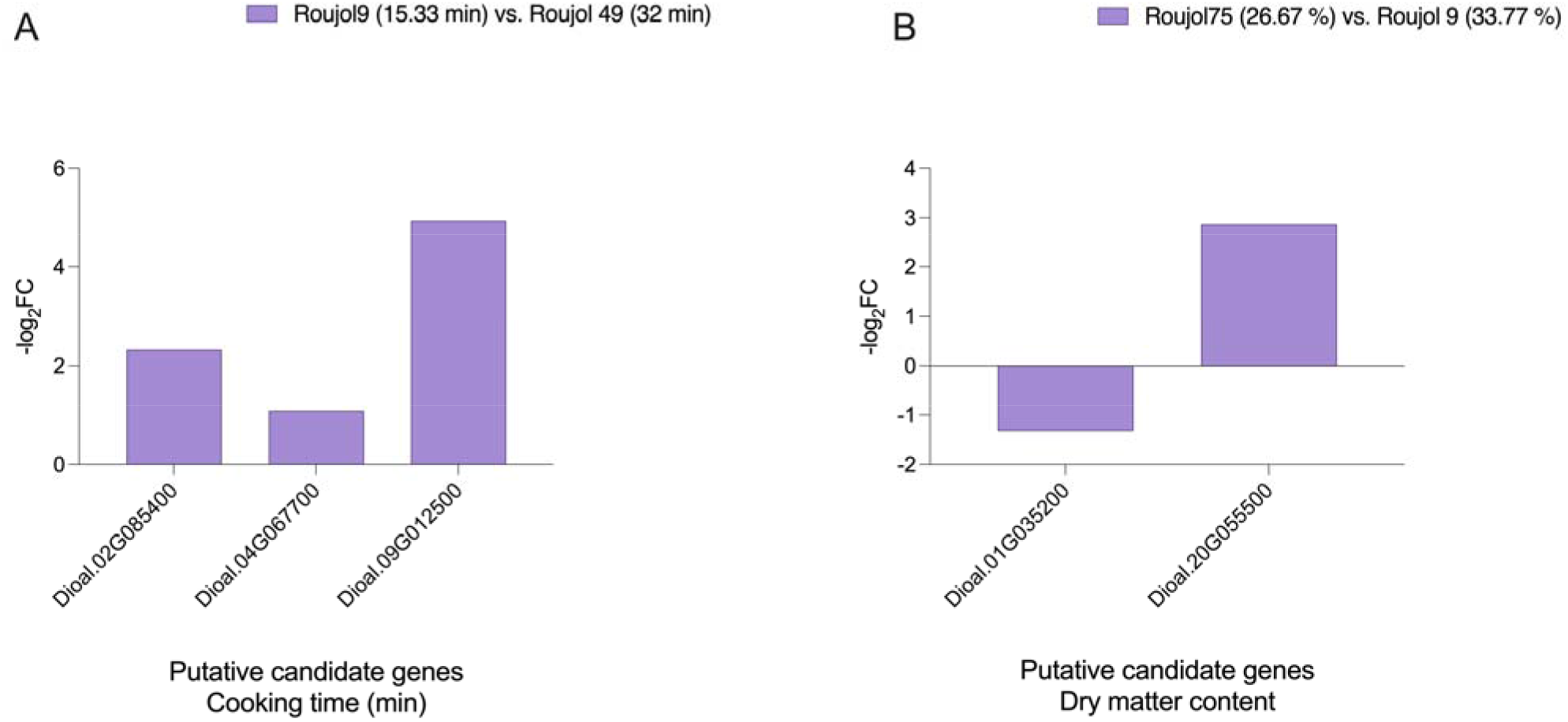
Differential expression of candidate genes identified in various comparisons of genotypes with contrasting culinary qualities. **A)** Expression profile of differentially expressed putative candidate genes for cooking time (min) in comparison Roujol9 vs. Roujol49 **B)** Differentially expressed putative candidate genes for dry matter contents in comparison Roujol75 vs. Roujol9. The comparisons were selected based on values in parenthesis for each trait.

Moreover, concerning dry matter content, two candidate genes, *Dioal.01G035200* and *Dioal.20G055500,* were identified with differential expression patterns in contrasting genotypes (Roujol75 vs. Roujol9). Roujol75 has a lower DM (26.67 %) compared to Roujol9 (33.77 %).

## Discussion

The culinary quality of yam is a crucial aspect that determines its consumers’ choices and various processing purposes [36]. Understanding and improving yam quality is of great importance to farmers, consumers, and the yam industry [25, 45]. It involves the identification of genetic factors, environmental influences, and processing techniques that can enhance desirable quality traits and ensure consistent product performance. Through scientific research, including genetic studies, sensory evaluations, and processing optimization, efforts are being made to enhance yam quality and promote its utilization as a nutritious staple worldwide [34, 35, 46]. In this study, we conducted a comprehensive evaluation of a diverse panel of *Dioscorea alata* genotypes to investigate the genetic basis of culinary attributes, including cooking time, dry matter content, hardness, and moldability. We employed genome-wide association studies (GWAS) to identify significant associations between genetic markers and the observed variations in these traits.

The observed phenotypic variation between Roujol and Godet in terms of quality traits in yam can be attributed to environmental influences. Previous studies showed that environmental factors significantly influence yam yield [47, 48], but multi-location trials assessing environmental impacts on quality traits are lacking [36]. Conducting multi-location trials across diverse environments is crucial to fully understand genotype-by-environment interactions and identify stable, high-quality genotypes [49]. Moreover, establishing trait correlations can reveal proxy markers to improve the prediction of yam quality [50]. For instance, correlating quality traits like dry matter content, starch composition, and texture profiles with sensory and cooking properties could reveal proxy traits that predict eating quality. Validating predictive quality models based on rapid analytical methods for proxies could enable robust multi-trait selection [51]. This offers a path to accelerated breeding gains for culinary quality without exhaustive cooking assays. Previous work by Ehounou et al. [52] has proposed that high protein content, low hardness, and high cohesiveness may contribute to reduced moldability in yam tubers. Based on these findings, they suggested using these criteria for selecting moldable genotypes in breeding programs. Supporting these associations, our observations, DM to be associated with improved moldability and increased hardness in yam tubers, align with previous root and tuber crops studies such as cassava [53], sweet potato [54], and potato [55]. Strong correlation and high heritability suggest DM as a robust proxy for accelerating indirect improvement of yam culinary quality, complemented by other potential proxies like starch and texture. Moreover, our diversity panel exhibits sufficient variation for multiple quality traits, enabling association mapping.

The presence of a high degree of recombination in *D. alata*, as suggested in a previous report [36], underscores the importance of utilizing high SNP density for GWAS in yam. We validated this by conducting GWAS using low-density (45,000 SNPs) and high-density (1.9 million SNPs) SNP datasets. Our findings confirmed that high SNP density significantly improves the GWAS results in *D. alata*. This high resolution allowed for more precise identification of associations between genetic markers and the studied traits. In addition, our results also demonstrated that the multi-locus FarmCPU model outperformed the single-locus MLM model in terms of identifying key associations between genetic markers and the studied traits. These findings are consistent with previous reports highlighting the superior performance of multi-locus GWAS models compared to single-locus models [56–58].

Food texture plays a crucial role as a quality indicator in various food products, including boiled and pounded yam [21]. It encompasses the structural and mechanical properties of the food, as well as the sensory perception experienced during handling and consumption. Parameters such as DM, fiber percentage, mealiness, waxiness, stickiness, and hardness are important textural attributes for boiled yam [59], which vary across different genotypes and environments [59, 60]. Bredeson et al. [35] identified a QTL on chromosome 18 associated with DM in *D. alata*. Gatarira et al. [41] studied DM in yam tubers at four locations but did not report stable SNP. In our study, we identified a stable SNP associated with DM on chromosome 9 (SNP rs839254). This SNP is located downstream of the *Dioal.09G047900* gene, which encodes a Winged helix-turn-helix DNA-binding domain protein. We propose that *Dioal.09G047900* may play a significant role for DM content in *D. alata*. Two differentially expressed genes identified between high and low dry matter genotypes encode a No Apical Meristem-associated C-terminal domain (NAM) transcription factor (*Dioal.01G035200*; SNP rs50615) and a protein kinase domain-containing protein (*Dioal.20G055500*; SNP rs1949352). NAM transcription factors regulate diverse growth, developmental, and stress response processes in plants [61], while protein kinases play critical roles in signaling cascades and cellular regulation [62]. For instance, a previous report found that overexpression of a calcium-dependent protein kinase in potato supported tuber wound healing [63]. However, neither of these genes has been functionally characterized in yam. Further investigation and molecular studies are warranted to elucidate the potential mechanisms by which these differentially expressed genes may modulate DM in *D. alata*. There are limited tools available currently to test candidate genes, and GWAS hits in yams [64]. For instance, CRISPR-Cas genome editing system could be very useful for improving yams through functional genomics [65, 66]. “Cut-dip-budding” delivery system [67] provides a way to deliver CRISPR-Cas to edit and study these genes without the need for transformation and regeneration and enables rapid, efficient CRISPR editing while minimizing off-target effects and eliminating transgenes.

Similarly, we identified a stable SNP for cooking time on chromosome 1. This particular SNP was located in an intergenic region between two genes, *Dioal.01G006400* and *Dioal.01G006500*. Of these two genes, *Dioal.01G006400* encodes a Tripeptidyl peptidase (TPP) known enzyme that plays an important role in protein breakdown and metabolism in plants [68, 69]. This could have implications for the breakdown of proteins during cooking, potentially affecting the cooking time of yam tubers by manipulating protein turnover [70]. Moreover, three genes, encoding Threonine protein kinase, ABL interactor-like protein 2, and Transposon protein, identified in the close proximity of GWAS associations were also identified as DEGs in two contrasting genotypes for cooking time.

The identification of *Dioal.04G099800*, which encodes a Steroid-transporting ATPase protein, in the vicinity of the association signal on chromosome 4 suggests its potential role in regulating moldability in *D. alata*. ABC transporters, including their ATP-binding proteins, are known for their involvement in the transport of various molecules across cellular membranes [71]. For example, in some root crops, the ABC transporters may be involved in the transport of starch, which is a major component contributing to the hardness or firmness of the roots [72]. These transporters could play a role in the uptake and storasage of starch in tubers, influencing the texture, moldability, and firmness of the tubers.

In this study, transcriptomic analysis was performed on mature tubers, which precluded the identification of DEGs during earlier developmental stages that may influence trait variation. Additionally, many genotypes shared the same alleles at peak SNPs identified through GWAS, limiting pairwise comparisons. To better elucidate DEGs and validate candidate genes, future studies could examine gene expression at multiple tuber developmental time-points using RNA-seq and qRT-PCR [73, 74]. Comparing genotypes with contrasting alleles at peak GWAS SNPs would help connect transcriptional profiles to sequence polymorphisms influencing the phenotype. Examining immature tubers would provide greater insight into transcriptional regulation during critical stages of tuber formation and expansion.

The GWAS peaks and putative candidate genes identified in this study represent novel findings in the context of *D. alata* and their association with quality attributes. These findings expand our understanding of the genetic basis of the culinary attributes of *D. alata*. Future studies should expand beyond core traits like dry matter content to explore additional biochemical factors affecting texture and taste, such as amylose content and pectin levels (https://rtbfoods.cirad.fr/). Transitioning significant marker-trait associations into KASP or SNP markers will facilitate breeding tasks [75]. Additionally, adopting high-throughput phenomic tools like NIRS [76, 77] modeling for tuber quality traits will accelerate selective breeding. Recent research demonstrates the potential of NIRS analysis on raw tubers or flours for rapid, nondestructive evaluation of breeding populations [7, 77–79]. Integrating NIRS-enabled phenotyping into selection pipelines could greatly increase genetic gains for culinary quality. By combining expanded genomics research, biotechnology advances like CRISPR editing, and front-end phenotypic screening, transformative improvements in yam culinary quality are within reach.

## Data availability

The raw sequencing data are available in the NCBI Sequence Read Archive under the BioProject number: PRJNA880983. The raw RNA-seq data are available under the BioProject number: PRJNA918625.

## Acknowledgments

We thank Angélique Morel, Marie-Claire Gravillon, Christophe Perrot, and Elie Nudol for their assistance during the experiments. We thank Ana Zotta Mota for her assistance in processing the RNA-seq data.

## Author Contributions

*Conceptualization*: Komivi Dossa, Denis Cornet. Investigation and data collection: Komivi Dossa, Mahugnon Ezékiel Houngbo, Mathieu Lechaudel, Erick Malédon, Jean-Luc Irep, *Data curation and Formal analysis*: Komivi Dossa, Mahugnon Ezékiel Houngbo, Yedomon Ange Bovys Zoclanclounon, Mian Faisal Nasir, Mathieu Lechaudel, Hâna Chair, Denis Cornet. *Funding acquisition*: Denis Cornet, Hâna Chair. *Writing ± original draft*: Komivi Dossa. *Writing ± review & editing*: Mathieu Lechaudel, Denis Cornet, Hâna Chair. All authors have read and approved the final version of this manuscript.

## Supplementary data

**Table S1.** Phenotypic charaterization of 52 genotypes of *Dioscorea alata*

**Table S2.** Selected genotypes for expression profiling of candidate genes for each trait under study

**Figure S1. Genome wide association studies for hardness and moldability A)** The Manhattan plot for hardness displays the genomic positions of SNPs and their association p-values specifically for the hardness, **B)** The QQ-plot associated with hardness depicts the observed p-values against the expected p-values under the null hypothesis of no association. Deviations from the diagonal line suggest potential associations between SNPs and hardness, **C)** The Manhattan plot for moldability at Godet displays the genomic positions of SNPs and their association p-values specifically for the hardness, **D)** The QQ-plot associated with moldability at Godet, **E)** The Manhattan plot for moldability at Roujol displays the genomic positions of SNPs and their association p-values specifically for the hardness, **F)** The QQ-plot associated with moldability at Roujol.

**Figure S2. The allele segregation analysis at peak SNPs A)** allele segregation analysis of rs1634685 demonstrates the relationship between different genotypes and their corresponding hardness. In this analysis, the TC allele is associated with genotypes that have increased hardness, while the TT allele is associated with genotypes that have lower hardness **B)** The allele segregation analysis concerning SNP rs381001 demonstrates the relationship between different genotypes and their corresponding moldability. In this analysis, the AA allele is associated with genotypes that have lower moldability, CC allele is associated with genotypes that having higher moldability, while the AC allele is associated with genotypes that have medium moldability.

**Figure S3.** Genome wide mapping of QTNs identified in current study and previously published QTLs identified for dry matter content in *Dioscorea alata*.

## References

1. Arnau G, Abraham K, Sheela M, Chair H, Sartie A, Asiedu R: Yams. Root and tuber crops 2010:127–148.

2. Headland TN: The wild yam question: How well could independent hunter-gatherers live in a tropical rain forest ecosystem? Human Ecology 1987:463–491.

3. Bahuchet S, McKey D, De Garine I: Wild yams revisited: Is independence from agriculture possible for rain forest hunter-gatherers? Human Ecology 1991, 19:213–243.

4. Omosowon S: The enantiophyllum clade of dioscorea in Africa: Systematics, distribution and conservation assessment. 2018.

5. Sharif BM, Burgarella C, Cormier F, Mournet P, Causse S, Van KN, Kaoh J, Rajaonah MT, Lakshan SR, Waki J: Genome-wide genotyping elucidates the geographical diversification and dispersal of the polyploid and clonally propagated yam (Dioscorea alata). Annals of botany 2020, 126(6):1029–1038.

6. Darkwa K, Olasanmi B, Asiedu R, Asfaw A: Review of empirical and emerging breeding methods and tools for yam (Dioscorea spp.) improvement: Status and prospects. Plant Breeding 2020, 139(3):474–497.

7. Ehounou AE, Cornet D, Desfontaines L, Marie-Magdeleine C, Maledon E, Nudol E, Beurier G, Rouan L, Brat P, Lechaudel M: Predicting quality, texture and chemical content of yam (Dioscorea alata L.) tubers using near infrared spectroscopy. Journal of Near Infrared Spectroscopy 2021, 29(3):128–139.

8. Kim D-S, Choi M-H, Shin H-J: Estimation of starch hydrolysis in sweet potato (Beni Haruka) based on storage period using nondestructive near-infrared spectrometry. Agriculture 2021, 11(2):135.

9. Alamu EO, Nuwamanya E, Cornet D, Meghar K, Adesokan M, Tran T, Belalcazar J, Desfontaines L, Davrieux F: Near-infrared spectroscopy applications for high-throughput phenotyping for cassava and yam: A review. International journal of food science & technology 2021, 56(3):1491–1501.

10. Tortoe C, Nketia S, Owusu M, Akonor PT, Dowuona S, Otoo E: Sensory attributes and consumer preference of precooked vacuum-packaged yam from two varieties of Ghanaian yam (Dioscorea rotundata) in the Accra metropolitan area. 2014.

11. Brunnschweiler J: Structure and texture of yam (Dioscorea spp.) and processed yam products. ETH Zurich; 2004.

12. Ebah D, Catherine B, Diby NNA, Kanon L, Oura R, Kouakou AM: State of knowledge on fresh yam and pounded yam in Côte d’Ivoire. Understanding the drivers of trait preferences and the development of multi-user RTB product profiles, WP1. 2023.

13. Alamu EO, Adesokan M, Awoyale W, Oyedele H, Fawole S, Asfaw A, Maziya-Dixon B: Assessment of biochemical, cooking, sensory and textural properties of the boiled food product of white yam (D. rotundata) genotypes grown at different locations. Heliyon 2022, 8(12):e11690.

14. Medoua GN, Mbome IL, Agbor-Egbe T, Mbofung C: Study of the hard-to-cook property of stored yam tubers (Dioscorea dumetorum) and some determining biochemical factors. Food research international 2005, 38(2):143–149.

15. Grundy MM-L, Edwards CH, Mackie AR, Gidley MJ, Butterworth PJ, Ellis PR: Re-evaluation of the mechanisms of dietary fibre and implications for macronutrient bioaccessibility, digestion and postprandial metabolism. British Journal of Nutrition 2016, 116(5):816–833.

16. Lai YC, Wang SY, Gao HY, Nguyen KM, Nguyen CH, Shih MC, Lin KH: Physicochemical properties of starches and expression and activity of starch biosynthesis-related genes in sweet potatoes. Food chemistry 2016, 199:556–564.

17. Sebio L, Chang Y: Effects of selected process parameters in extrusion of yam flour (Dioscorea rotundata) on physicochemical properties of the extrudates. Food/Nahrung 2000, 44(2):96–101.

18. Babajide J, Henshaw F, Oyewole O: EFFECT OF YAM VARIETY ON THE PASTING PROPERTIES AND SENSORY ATTRIBUTES OF TRADITIONAL DRY-YAM AND ITS PRODUCTS. Journal of Food Quality 2008, 31(3):295–305.

19. Otegbayo B, Aina J, Abbey L, SAKYI-DAWSON E, Bokanga M, Asiedu R: Texture profile analysis applied to pounded yam. Journal of texture studies 2007, 38(3):355–372.

20. Awoyale W, Olatoye KK, Maziya-Dixon B: Cassava Pectin and Textural Attributes of Cooked gari (eba) and fufu Dough. In: Utilization of Pectin in the Food and Drug Industries. IntechOpen; 2023.

21. Otegbayo B, Madu T, Oroniran O, Chijioke U, Fawehinmi O, Okoye B, Tanimola A, Adebola P, Obidiegwu J: End-user preferences for pounded yam and implications for food product profile development. International Journal of Food Science & Technology 2021, 56(3):1458–1472.

22. Tran T, Zhang X, Ceballos H, Moreno JL, Luna J, Escobar A, Morante N, Belalcazar J, Becerra LA, Dufour D: Correlation of cooking time with water absorption and changes in relative density during boiling of cassava roots. International journal of food science & technology 2021, 56(3):1193–1205.

23. Safo-Kantanka O, Owusu-Nipah J: Cassava varietal screening for cooking quality: relationship between dry matter, starch content, mealiness and certain microscopic observations of the raw and cooked tuber. Journal of the Science of Food and Agriculture 1992, 60(1):99–104.

24. Lebot V, Prana MS, Kreike N, Van Heck H, Pardales J, Okpul T, Gendua T, Thongjiem M, Hue H, Viet N: Characterisation of taro (Colocasia esculenta (L.) Schott) genetic resources in Southeast Asia and Oceania. Genetic Resources and Crop Evolution 2004, 51:381–392.

25. Asfaw A, Agre P, Matsumoto R, Olatunji AA, Edemodu A, Olusola T, Odom-Kolombia OL, Adesokan M, Alamu OE, Adebola P: Genome-wide dissection of the genetic factors underlying food quality in boiled and pounded white Guinea yam. Journal of the Science of Food and Agriculture 2023.

26. Khakasa E, Muyanja C, Mugabi R, Bugaud C, Forestier-Chiron N, Uwimana B, Arinaitwea IK, Nowakundaa K: Sensory characterization of the perceived quality of East African highland cooking. 2023.

27. Osunbade AO, Alamu EO, Awoyale W, Akinwande AB, Adejuyitan JA, Maziya-Dixon B: End-user quality characteristics and preferences for cassava, yam and banana products in rural and urban areas-A review. Cogent Food & Agriculture 2023, 9(1):2205720.

28. Ufondu HE, Maziya-Dixon B, Okoyeuzu CF, Okonkwo TM, Okpala COR: Effects of yam varieties on flour physicochemical characteristics and resultant instant fufu pasting and sensory attributes. Scientific Reports 2022, 12(1):20276.

29. Obidiegwu JE, Lyons JB, Chilaka CA: The Dioscorea Genus (Yam)—An appraisal of nutritional and therapeutic potentials. Foods 2020, 9(9):1304.

30. Mignouna H, Mank R, Ellis T, Van Den Bosch N, Asiedu R, Abang M, Peleman J: A genetic linkage map of water yam (Dioscorea alata L.) based on AFLP markers and QTL analysis for anthracnose resistance. Theoretical and Applied Genetics 2002, 105:726–735.

31. Cornet D, Sierra J, Tournebize R, Gabrielle B, Lewis FI: Bayesian network modeling of early growth stages explains yam interplant yield variability and allows for agronomic improvements in West Africa. European journal of agronomy 2016, 75:80–88.

32. Dansi A, Dantsey-Barry H, Dossou-Aminon I, N’kpenu E, Agré A, Sunu Y, Kombaté K, Loko Y, Dansi M, Assogba P: Varietal diversity and genetic erosion of cultivated yams (Dioscorea cayenensis Poir-D. rotundata Lam complex and D. alata L.) in Togo. Int J Biodivers Conserv 2013, 5(2):223–239.

33. Mondo JM, Agre PA, Edemodu A, Asiedu R, Akoroda MO, Asfaw A: Cross compatibility in intraspecific and interspecific hybridization in yam (Dioscorea spp.). Scientific reports 2022, 12(1):3432.

34. Ehounou AE, Cormier F, Maledon E, Nudol E, Vignes H, Gravillon MC, N’guetta ASP, Mournet P, Chaïr H, Kouakou AM: Identification and validation of QTLs for tuber quality related traits in greater yam (Dioscorea alata L.). Scientific Reports 2022, 12(1):1–14.

35. Bredeson JV, Lyons JB, Oniyinde IO, Okereke NR, Kolade O, Nnabue I, Nwadili CO, Hřibová E, Parker M, Nwogha J: Chromosome evolution and the genetic basis of agronomically important traits in greater yam. Nature communications 2022, 13(1):1–16.

36. Dossa K, Morel A, Houngbo ME, Mota AZ, Maledon E, Irep JL, Diman JL, Mournet P, Causse S, Van KN: Genome-wide association studies reveal novel loci controlling tuber flesh color and oxidative browning in Dioscorea alata. Journal of the Science of Food and Agriculture 2023.

37. Mota AZ, Dossa K, Lechaudel M, Cornet D, Mournet P, Lopez D, Chaïr H: Genomic insights into greater yam tuber quality traits. bioRxiv 2023:2023.2003. 2017.532727.

38. Lu R-S, Hu K, Zhang F-J, Sun X-Q, Chen M, Zhang Y-M: Pan-plastome of greater yam (Dioscorea alata) in China: intraspecific genetic variation, comparative genomics, and phylogenetic analyses. International Journal of Molecular Sciences 2023, 24(4):3341.

39. Ehounou AE, Cormier F, Maledon E, Nudol E, Vignes H, Gravillon MC, N’guetta ASP, Mournet P, Chaïr H, Kouakou AM: Identification and validation of QTLs for tuber quality related traits in greater yam (Dioscorea alata L.). Scientific Reports 2022, 12(1):8423.

40. Bredeson JV, Lyons JB, Oniyinde IO, Okereke NR, Kolade O, Nnabue I, Nwadili CO, Hřibová E, Parker M, Nwogha J: Chromosome evolution and the genetic basis of agronomically important traits in greater yam. Nature communications 2022, 13(1):2001.

41. Gatarira C, Agre P, Matsumoto R, Edemodu A, Adetimirin V, Bhattacharjee R, Asiedu R, Asfaw A: Genome-wide association analysis for tuber dry matter and oxidative browning in water yam (Dioscorea alata L.). Plants 2020, 9(8):969.

42. Mondo JM, Agre PA, Asiedu R, Akoroda MO, Asfaw A: Genome-wide association studies for sex determination and cross-compatibility in water yam (Dioscorea alata L.). Plants 2021, 10(7):1412.

43. Arnau G, Bhattacharjee R, Mn S, Chair H, Malapa R, Lebot V, Perrier X, Petro D, Penet L, Pavis C: Understanding the genetic diversity and population structure of yam (Dioscorea alata L.) using microsatellite markers. PLoS One 2017, 12(3):e0174150.

44. Yin L: CMplot: circle manhattan plot. R package version 2020, 3(2).

45. Akissoe N, Mestres C, Hounhouigan J, Nago M: Biochemical origin of browning during the processing of fresh yam (Dioscorea spp.) into dried product. Journal of Agricultural and Food Chemistry 2005, 53(7):2552–2557.

46. Ray DK, Ramankutty N, Mueller ND, West PC, Foley JA: Recent patterns of crop yield growth and stagnation. Nature communications 2012, 3(1):1–7.

47. Asfaw A: Standard operating protocol for yam variety performance evaluation trial. In.; 2016.

48. Norman PE, Tongoona PB, Danquah A, Danquah EY, Agre PA, Agbona A, Asiedu R, Asfaw A: Genetic Analysis of Agronomic and Quality Traits from Multi-Location white Yam Trials using Mixed Model with Genomic Relationship Matrix. Global Journal of Botanical Science 2022, 10:8–22.

49. Crossa J: Statistical analyses of multilocation trials. Advances in agronomy 1990, 44:55–85.

50. Wasson AP, Richards R, Chatrath R, Misra S, Prasad SS, Rebetzke G, Kirkegaard J, Christopher J, Watt M: Traits and selection strategies to improve root systems and water uptake in water-limited wheat crops. Journal of experimental botany 2012, 63(9):3485–3498.

51. Mathai N, Chen Y, Kirchmair J: Validation strategies for target prediction methods. Briefings in bioinformatics 2020, 21(3):791–802.

52. Ehounou AE, Cormier F, Cornet D, Maledon E, Desfontaines L, Marie-Magdeleine C, Nudol E, Gravillon M-C, Kouakou AM, Arnau G: Development of NIRS and molecular marker to improve breeding efficiency in Greater Yam (Dioscorea alata L.) for key quality traits. ABB1103. In: 2019.

53. Egesi CN, Ilona P, Ogbe F, Akoroda M, Dixon A: Genetic variation and genotype× environment interaction for yield and other agronomic traits in cassava in Nigeria. Agronomy journal 2007, 99(4):1137–1142.

54. Aina AJ, Falade KO, Akingbala JO, Titus P: Physicochemical properties of twenty-one Caribbean sweet potato cultivars. International journal of food science & technology 2009, 44(9):1696–1704.

55. Li B, Xu X, Zhang L, Han J, Bian C, Li G, Liu J, Jin L: Above-ground biomass estimation and yield prediction in potato by using UAV-based RGB and hyperspectral imaging. ISPRS Journal of Photogrammetry and Remote Sensing 2020, 162:161–172.

56. Liu X, Huang M, Fan B, Buckler ES, Zhang Z: Iterative usage of fixed and random effect models for powerful and efficient genome-wide association studies. PLoS genetics 2016, 12(2):e1005767.

57. Zhong H, Liu S, Meng X, Sun T, Deng Y, Kong W, Peng Z, Li Y: Uncovering the genetic mechanisms regulating panicle architecture in rice with GPWAS and GWAS. BMC genomics 2021, 22:1–13.

58. Nazir MF, He S, Ahmed H, Sarfraz Z, Jia Y, Li H, Sun G, Iqbal MS, Pan Z, Du X: Genomic insight into the divergence and adaptive potential of a forgotten landrace G. hirsutum L. purpurascens. Journal of Genetics and Genomics 2021, 48(6):473–484.

59. Otegbayo B, Oroniran O, Fawehinmi O, Ayandiji A: State of Knowledge on Boiled and Pounded Yam in Nigeria.

60. Ezeocha V, Nwankwo I, Ezebuiro V: Evaluation of the chemical, functional and sensory properties of pre-release White Yam (Dioscorea rotundata) Genotypes in Umudike, Southeast, Nigeria. British Biotechnology Journal 2015, 9(4):1.

61. Zhang Y, Li D, Wang Y, Zhou R, Wang L, Zhang Y, Yu J, Gong H, You J, Zhang X: Genome-wide identification and comprehensive analysis of the NAC transcription factor family in Sesamum indicum. PloS one 2018, 13(6):e0199262.

62. Chen X, Ding Y, Yang Y, Song C, Wang B, Yang S, Guo Y, Gong Z: Protein kinases in plant responses to drought, salt, and cold stress. Journal of integrative plant biology 2021, 63(1):53–78.

63. Ma L, Jiang H, Ren Y-Y, Yang J-W, Han Y, Si H-J, Prusky D, Bi Y, Wang Y: Overexpression of StCDPK23 promotes wound healing of potato tubers by regulating StRbohs. Plant Physiology and Biochemistry 2022, 185:279–289.

64. Syombua ED, Zhang Z, Tripathi JN, Ntui VO, Kang M, George OO, Edward NK, Wang K, Yang B, Tripathi L: A CRISPR/Cas9-based genome-editing system for yam (Dioscorea spp.). Plant Biotechnology Journal 2021, 19(4):645.

65. Feng S, Song W, Fu R, Zhang H, Xu A, Li J: Application of the CRISPR/Cas9 system in Dioscorea zingiberensis. Plant Cell, Tissue and Organ Culture (PCTOC) 2018, 135:133–141.

66. Zaman QU, Raza A, Gill RA, Hussain MA, Wang HF, Varshney RK: New possibilities for trait improvement via mobile CRISPR-RNA. Trends in Biotechnology 2023.

67. Cao X, Xie H, Song M, Lu J, Ma P, Huang B, Wang M, Tian Y, Chen F, Peng J: Cut–dip– budding delivery system enables genetic modifications in plants without tissue culture. The Innovation 2023, 4(1).

68. Arima K, Uchikoba T, Yonezawa H, Shimada M, Kaneda M: Cucumisin-like protease from the latex of Euphorbia supina. Phytochemistry 2000, 53(6):639–644.

69. Silva-López R, Gonçalves R: Therapeutic proteases from plants: biopharmaceuticals with multiple applications. J Appl Biotechnol Bioeng 2019, 6(2):101–109.

70. Tomkinson B, Lindås A-C: Tripeptidyl-peptidase II: a multi-purpose peptidase. The international journal of biochemistry & cell biology 2005, 37(10):1933–1937.

71. Yazaki K: ABC transporters involved in the transport of plant secondary metabolites. FEBS letters 2006, 580(4):1183–1191.

72. Singh S, Selvakumar R, Mangal M, Kalia P: Breeding and genomic investigations for quality and nutraceutical traits in vegetable crops-a review. Indian Journal of Horticulture 2020, 77(1):1–40.

73. Cao T, Wang S, Ali A, Shan N, Sun J, et al: Transcriptome and metabolome analysis reveals the potential mechanism of tuber dynamic development in yam (*Dioscorea polystachya* Turcz.). LWT, 2023, 181, 114764. 10.1016/j.lwt.2023.114764.

74. Riekötter J, Oklestkova J, Muth J, Twyman RM, Epping J: Transcriptomic analysis of Chinese yam (*Dioscorea polystachya* Turcz.) variants indicates brassinosteroid involvement in tuber development. Front. Nutr. 2023, 10:1112793. doi: 10.3389/fnut.2023.1112793.

75. Cormier F, Martin G, Vignes H, Lachman L, Cornet D, Faure Y, Maledon E, Mournet P, Arnau G: Genetic control of flowering in greater yam (Dioscorea alata L.). BMC plant biology 2021, 21(1):1–12.

76. Van Tassel DL, DeHaan LR, Diaz-Garcia L, Hershberger J, Rubin MJ, Schlautman B, Turner K, Miller AJ: Re-imagining crop domestication in the era of high throughput phenomics. Current Opinion in Plant Biology 2022, 65:102150.

77. Lebot V, Malapa R: Application of near infrared reflectance spectroscopy for the evaluation of yam (Dioscorea alata) germplasm and breeding lines. Journal of the Science of Food and Agriculture 2013, 93(7):1788–1797.

78. Houngbo ME, Desfontaines L, Diman JL, Arnau G, Mestres C, Davrieux F, Rouan L, Beurier G, Carine MM, Meghar K: Convolutional neural network allows amylose content prediction in yam (Dioscorea alata L.) flour using near infrared spectroscopy. Journal of the Science of Food and Agriculture 2023.

79. Otegbayo B, Oroniran O, Tanimola A, Fawehinmi B, Alamu A, Bolaji T, Madu T, Okoye B, Chijioke U, Ofoeze M: Food quality profile of pounded yam and implications for yam breeding. Journal of the Science of Food and Agriculture 2023.

