## Supplementary figures and images for "Detecting the genetic variants associated with key culinary traits in *Dioscorea alata*"

### Figure S1

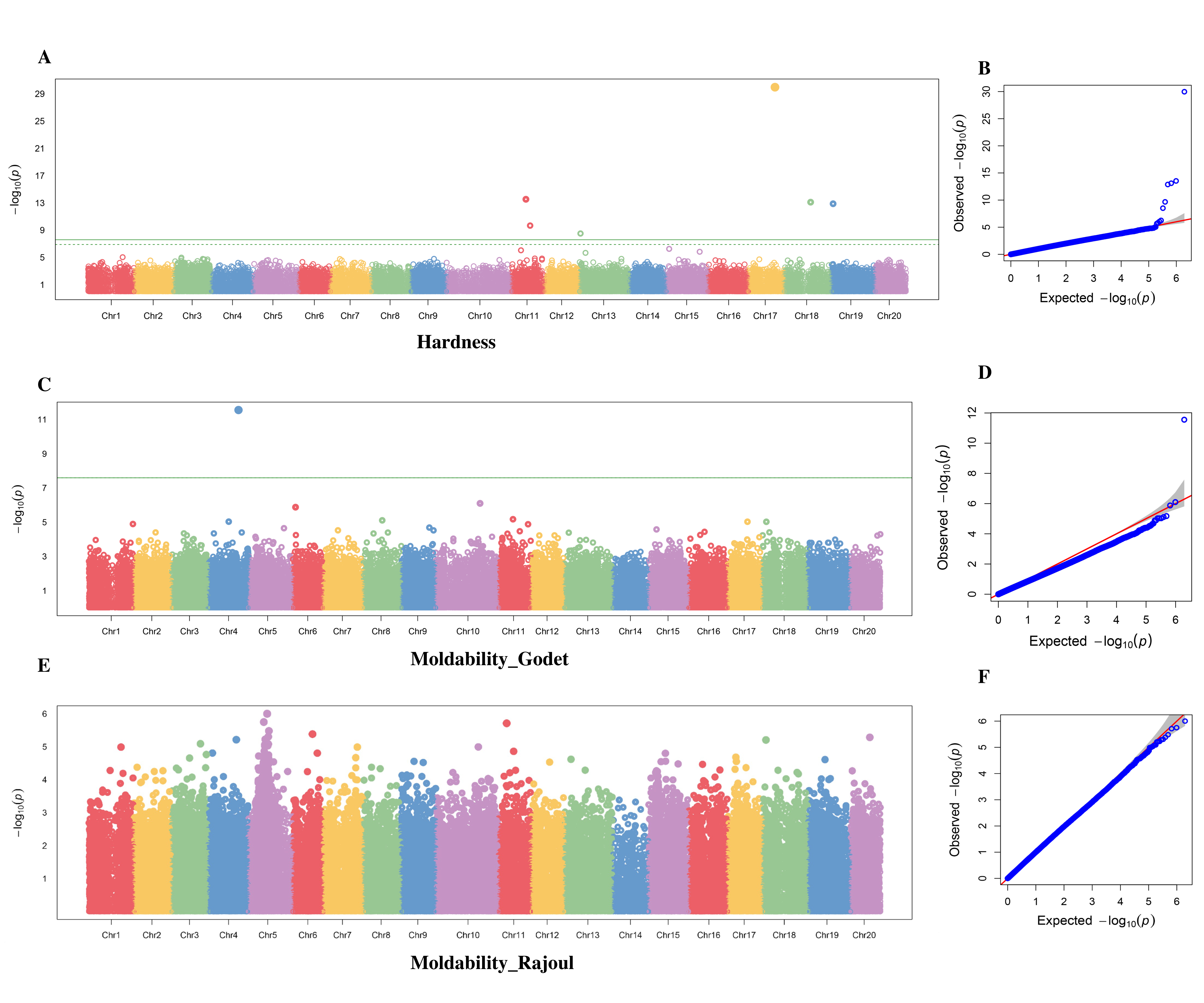

### Figure S2

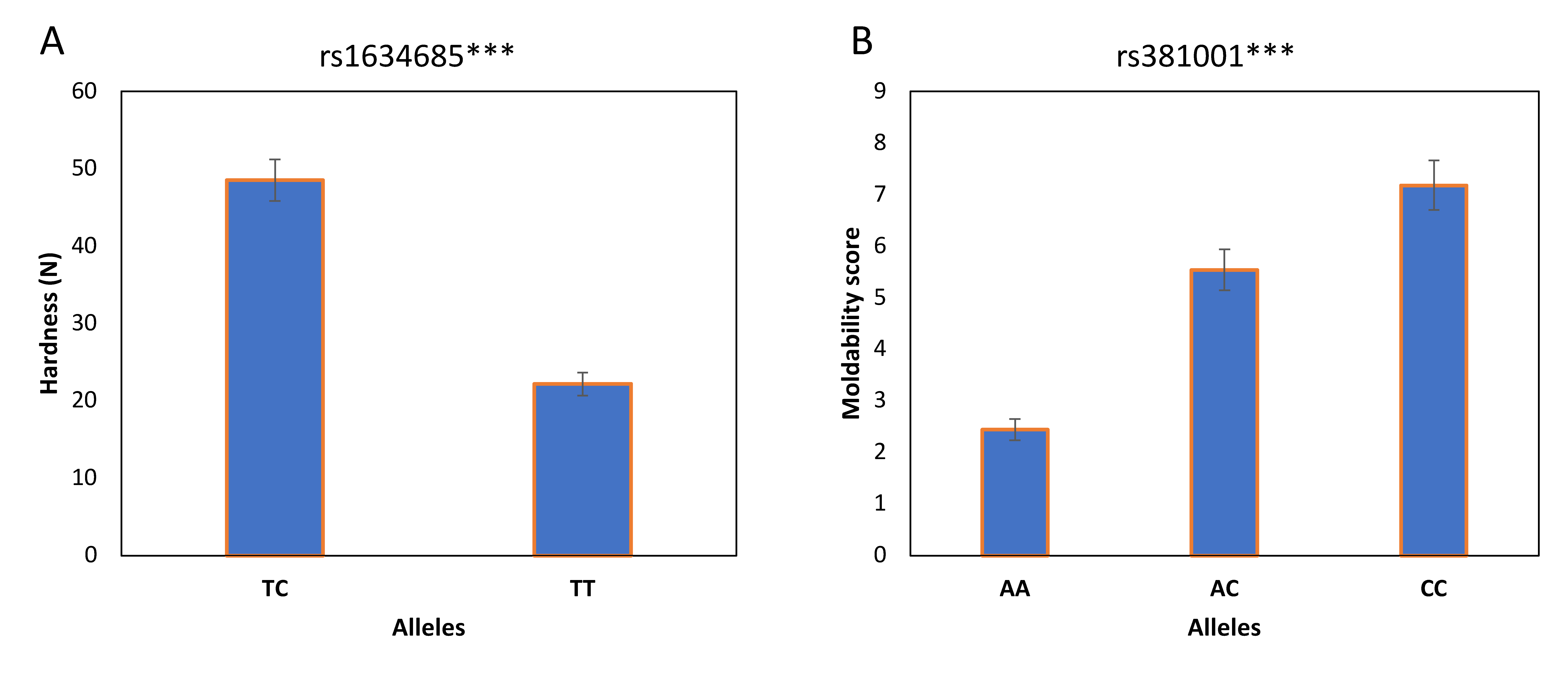

### Figure S3

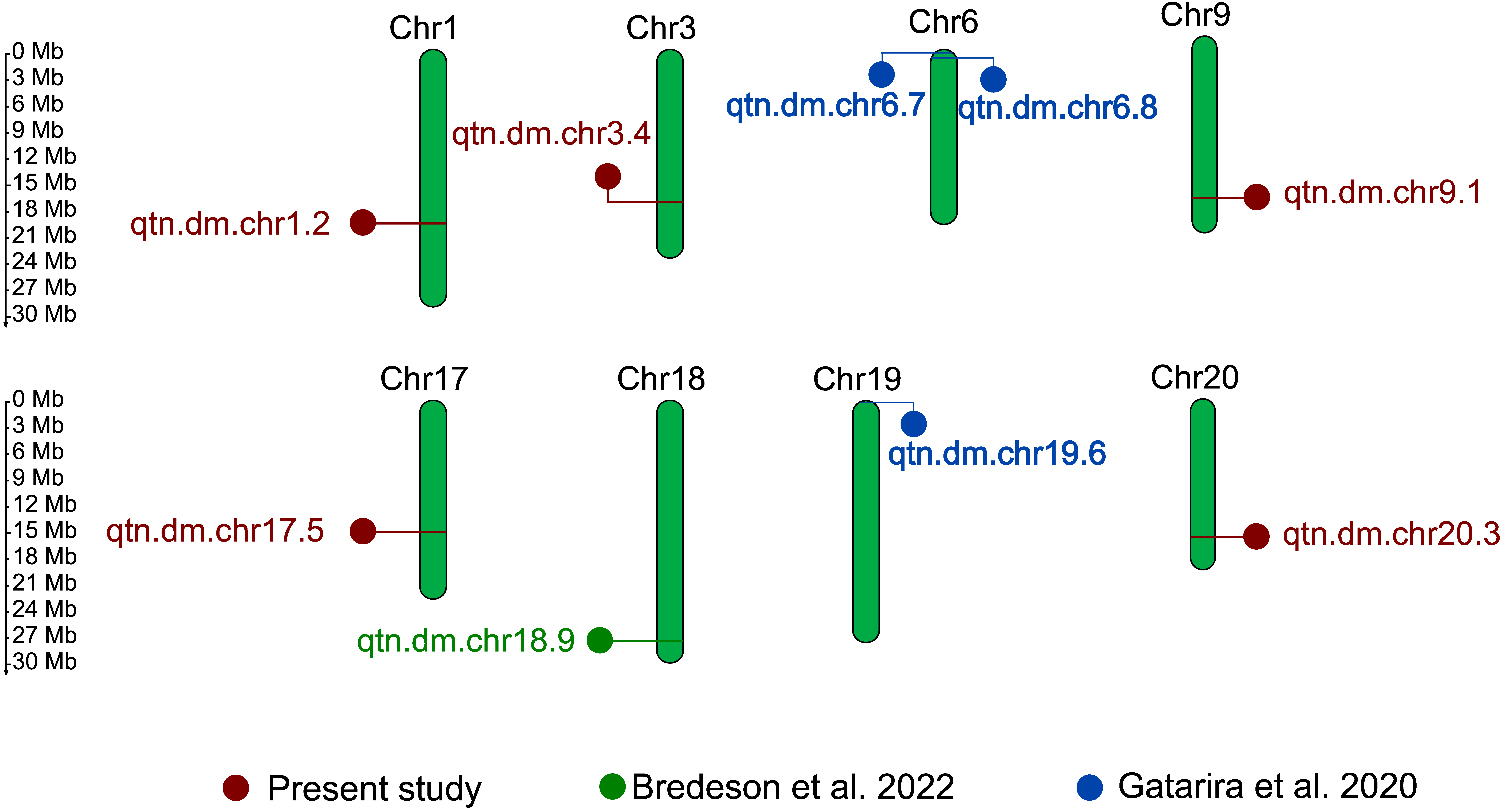
